# Axons of cortical basket cells originating from dendrites develop higher local complexity than axons emerging from basket cell somata

**DOI:** 10.1101/2023.08.28.555080

**Authors:** Steffen Gonda, Christian Riedel, Andreas Reiner, Ina Köhler, Petra Wahle

## Abstract

Neuronal differentiation is regulated by neuronal activity. Here, we analyzed dendritic and axonal growth of Basket cells and non-Basket cells using sparse transfection of channelrhodopsin-YFP and repetitive optogenetic stimulation in slice cultures of rat visual cortex. Neocortical interneurons often display axon-carrying dendrites (AcDs). We found that the AcDs of Basket cells and non-Basket cells were on average their most complex dendrite. Further, the AcD configuration had an influence on Basket cell axonal development. Axons originating from an AcD formed denser arborizations with more terminal endings within the dendritic field of the parent cell. Intriguingly, this occurred already in unstimulated Basket cells, and complexity was not increased further by optogenetic stimulation. However, optogenetic stimulation exerted a growth-promoting effect on axons emerging from Basket cell somata. The axons of non-Basket cells neither responded to the AcD configuration nor to the optogenetic stimulation. The results suggest that the formation of locally dense Basket cell plexuses is regulated by spontaneous activity. Moreover, in the AcD configuration, the AcD and the axon it carries mutually support each others growth.

**Teaser:** Textbooks usually draw cortical neurons with axons emerging from the cell body. Axons emerging from dendrites have been considered a caprice of nature until recent work demonstrated that axon-carrying dendrites are neither a rare nor a random phenomenon. Rather, this morphological configuration awards specific functional properties. Here, we discovered a morphogenetic role of the AcD configuration in developing Basket neurons which deliver axo-somatic inhibition to pyramidal neurons. Driven by spontaneous activity, AcD and the axon mutually promote each others growth such that the axon rapidly forms a denser presynaptic terminal field. Basket cells with axons from the cell body achieve the same complexity with technical assistance via an optogenetic stimulation. This way, a dendritic origin accelerates Basket cell axonal development.

## Introduction

The cortical interneuron types are classified by distinct axonal projection and termination patterns (Meyer and Ferres-Torres, 1984). The type-specific features are created by transcription factors (Favuzzi et al., 2019), and genetic alterations evoke morphofunctional abnormalities. For instance, in fast-spiking Basket cells (BC), mTORC1 haploinsufficiency accelerates the development of axon terminal elements which is followed by a loss of perisomatic innervation in the adult, suggesting a role of mTOR for maintenance of parvalbumin cell connectivity (Amegandjin et al., 2021). Sox6 is required for parvalbumin bouton growth and stability (Munguba et al., 2021). Lack of MeCP2 accelerates development of parvalbumin cells in vivo as well as in organotypic culture (OTC) (Patrizi et al., 2020). Yet, genetic determination does not shield interneurons from environmental influences, in particular neuronal activity. For visual cortical interneurons, ablating the retinal input causes a loss of dendritic spines and synapses (Keck et al., 2011). Overexpression of GluA1(Q)flip and GluK2 glutamate receptor subunits increases dendritic complexity and spine density of interneurons (Hamad et al., 2011; Jack et al., 2019). Axons of cortical interneurons silenced via activation of hM4Di designer receptor display the same degree of complexity and bouton size as mock-stimulated neurons, but have less bouton terminaux, a proxy of presynapses (Gasterstädt et al., 2022). Silenced cortical BC retain a normal complexity but innervate a lower number of pyramidal cells presumably due to enhanced pruning of terminal elements (Chattopadhyaya et al., 2004; Di Cristo et al., 2004; Baho and Di Cristo, 2012). Reducing the GABA production and the amount of GABA released from BC impairs their ability to stabilize axo-somatic synapses (Chattopadhyaya et al., 2007). In BC, inflammatory cytokine-evoked hyperexcitability lowers GAD, Kv3.2 and synaptotagmin-2 expression, evokes calcium events at higher frequencies but lower amplitude, and while the axonal complexity remains unchanged, the number and the size of axonal boutons declines (Engelhardt et al., 2018). Further, axonal geometry of P12-18 hippocampal chandelier cells is shaped through remodeling (Steinecke et al., 2017), and axo-axonic boutons decline in number when activity increases because the chandelier cell action at the axon initial segment is depolarizing at this age (Pan-Vasquez et al., 2020). Of the non-fast-spiking, non-Basket cell (non-BC), reducing excitability via expression of Kir2.1 channels from late fetal to P8 reduces axonal complexity of calretinin and reelin neurons derived from the caudal ganglionic eminence (De Marco Garcia et al., 2011).

The ability to reorganize BC axons persists in the adult. Within hours of sensory deprivation by whisker plucking the horizontal axons of deprived barrel BC discard local collaterals and sprout into non-deprived barrels almost doubling their length (Marik et al., 2010). Prelimbic parvalbumin cells grow more complex axons upon chemogenetic stimulation (Stedehouder et al., 2018). Further, epilepsy has been found to decrease dentate gyrus axo-axonic synapse density while the density of axo-somatic synapses increases (Alhourani et al., 2020).

Together, the evidence supports the view that the morphological development of interneuronal axons is regulated by activity. The activity-dependent regulation of terminal elements and boutons has been well characterized in slice cultures at week 3-4, which revealed the importance of GABA synthesis for terminal element development (Chattopadhyaya et al., 2004; Di Cristo et al., 2004; Chattopadhyaya et al., 2007; Baho and Di Cristo, 2012). However, the potential impact of a cell’s activity on the total dimensions of the axon plexus was hardly ever assessed. Interneuronal axons frequently emerge from dendrites (Wahle et al., 2022). Axon-carrying dendrites (AcD) of hippocampal pyramidal neurons have been reported to be privileged with a higher excitability and the ability to elicit action potential firing which escapes somatic inhibition, also their AcD is longer than regular dendrites (Thome et al., 2014; Hodapp et al., 2022). Yet, it is unknown if the AcD configuration has a morphogenetic impact on interneuronal dendrites or axons. Here, we report that BC axons, but not non-BC axons originating from an AcD formed locally denser arborizations with more terminal endings even without optogenetic stimulation. In contrast, channelrodopsin-mediated optogenetic stimulation increased the local axonal density of BC axons of somatic origin. These results suggest that the development of local BC plexuses is promoted by spontaneous activity and that the AcD configuration has a morphogenetic role for developing dendrites and axons.

## Results

### Cell type-specific effects of activity on dendritic complexity

Roller tube cultures are spontaneously active, preserve the morphofunctional properties of the interneuron types, and allow reconstruction of complete axons (Klostermann and Wahle, 1999; Di Cristo et al., 2004; Chattopadhyaya et al., 2007; Baho and Di Cristo, 2012; Feldmeyer et al., 2018). We reconstructed axons of ChR2-eYFP labeled BC (Figure 1A) and non-BC with arcade (Figure 1B) and bitufted morphology relying on classical axonal features (Meyer and Ferres-Torres, 1984; Klostermann and Wahle, 1999; Staiger et al., 2015; Markram et al., 2015; Feldmeyer et al., 2018) (Figure 1 and Figure 1-figure supplement 1; see inclusion criteria in Material and methods). In rodent cortex *in vivo*, the cell types become recognizable around P10. Analysis of BC axon arborization in developing mouse cortex found that many presynapses are already close to their somatic targets at P14 (Micheva et al., 2021). Further, miniature inhibitory postsynaptic current (IPSC) frequency increases from P6-P15 in mouse somatosensory cortex pyramidal cells, with most IPSC originating from perisomatic synapses (Kobayashi et al 2008; Micheva et al., 2021). Indeed, neurons at DIV 15 in OTC were well differentiated with BC forming thick main axons giving rise to thin side branches with irregular-sized boutons (Figure 1C), terminal elements contacting somata (Figure 1D), bouton terminaux which are *bona fide* presynapses (Figure 1E_1_). Growth cones suggested that axons of BC and non-BC were actively growing at DIV 15 (Figure 1E_2_). Accordingly, optogenetic stimulation was applied from DIV 11-15, which overlaps the main growth phase of BC.

**Figure 1 with Figure 1 supplement 1.**
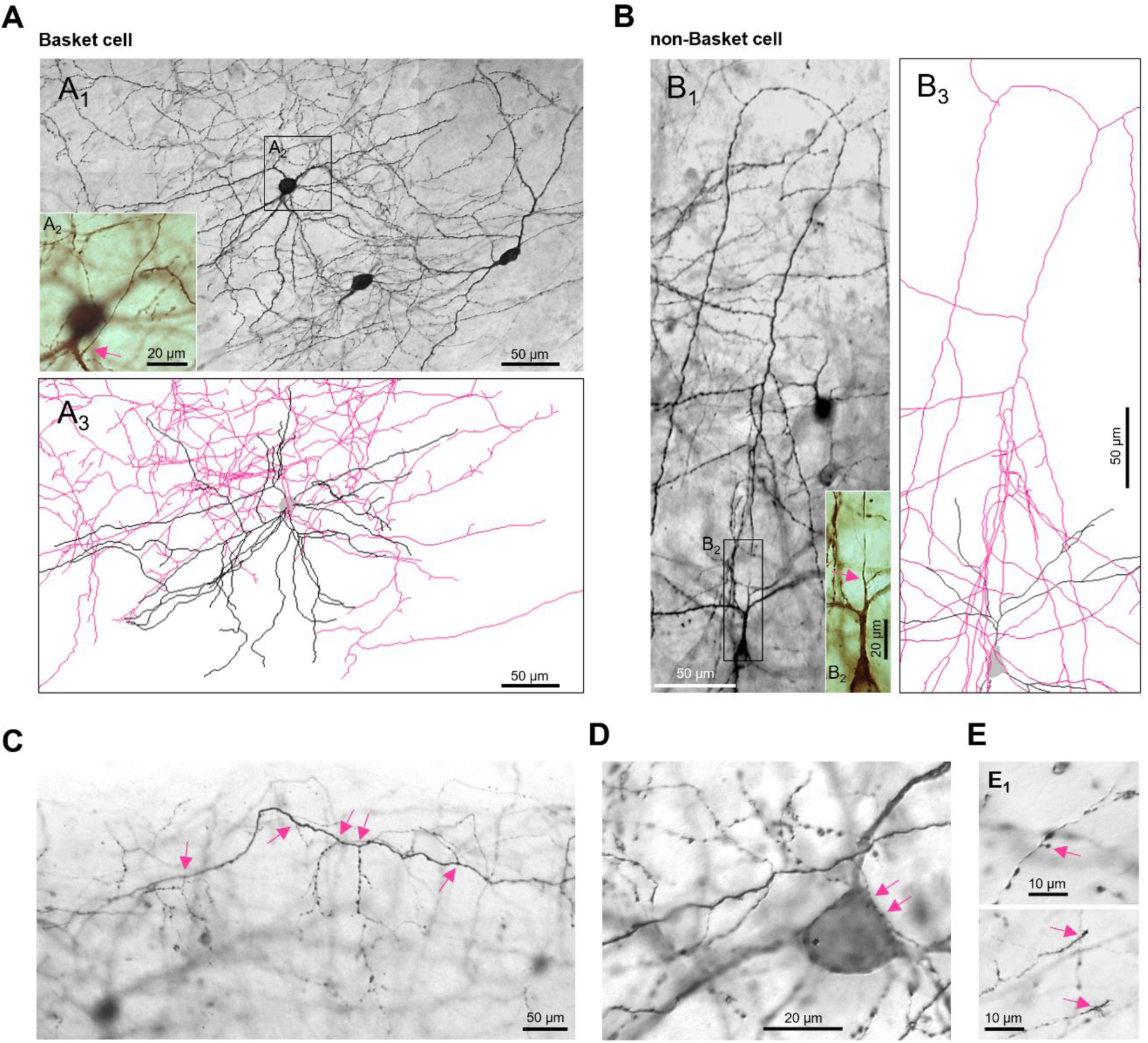
Representative interneurons with AcDs. (**A**) Basket cell; montage of 8 photomicrographs. A1. Cell overview. A2. The axon emerged from a dendrite close to the soma. A3. The local arborization, cropped from the Neurolucida reconstruction of the neuron. (**B**) Neuron with an arcade axon; montage of 3 photomicrographs. B1. The initial axonal branching. B2. The axon emerged from a second-order dendrite. B3. The local arborization, cropped from the Neurolucida reconstruction of the neuron. Immunohistochemical staining for transfected ChR2-YFP, diaminobenzidine reaction product intensified with osmium tetroxide for manual Neurolucida reconstruction at 1000x magnification. Axons in pink, dendrites in black, somata in gray. Pink arrows indicate the axon hillock. (**C**) Photomicrograph of a horizontal axon of a BC with local branches and termina elements; branch points indicated by arrows; the parent soma is in the background at the lower left corner. (**D**) Thick main axon of another BC giving rinse to a terminal branch contacting a pyramidal-shaped soma (arrows). (**E**) Axon collateral with a bouton terminaux (E1) and collaterals with growth cones at the tips (E2).

**Figure 1 – figure supplement 1.**
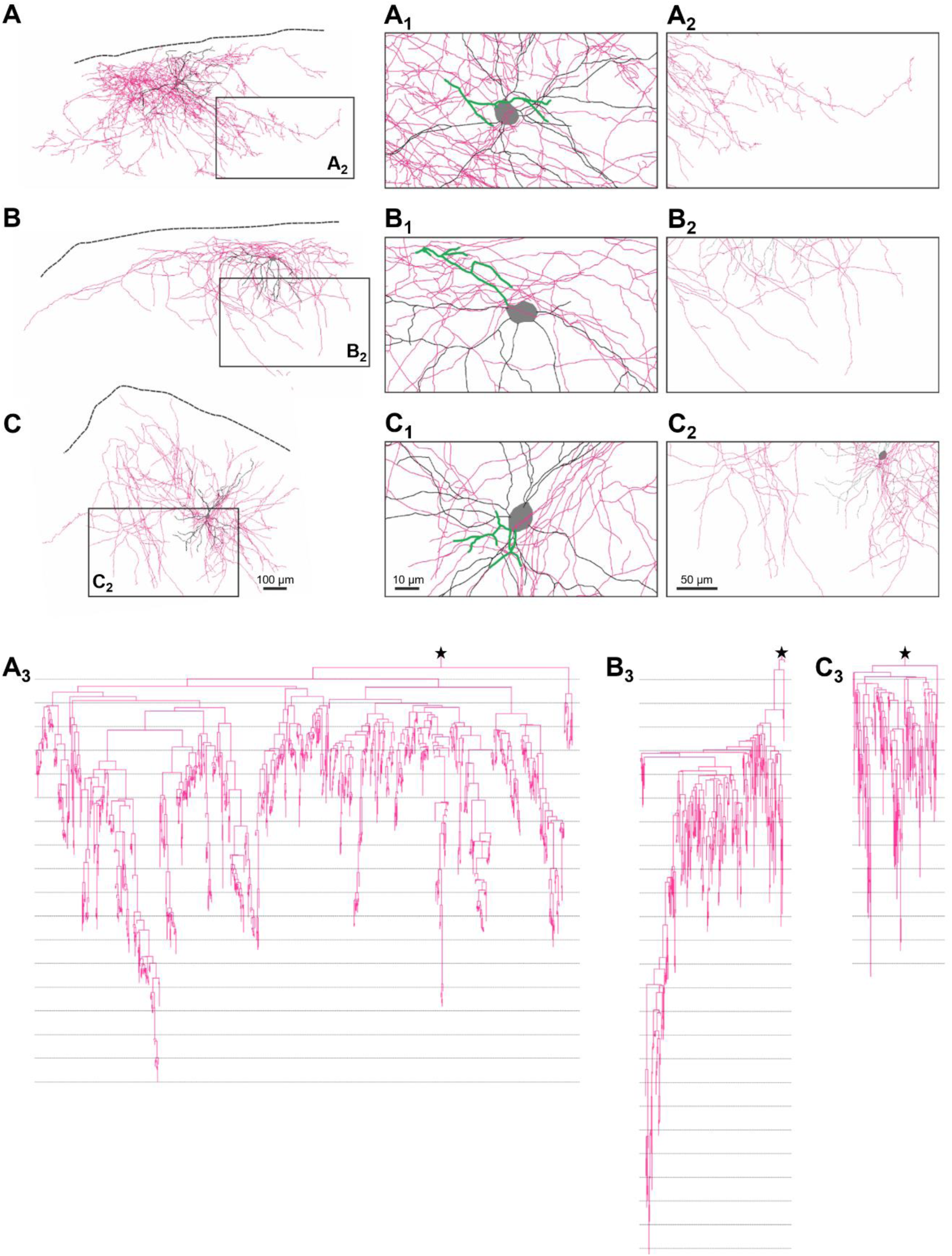
Reconstructions of representative neurons. (**A, A1-A3**) Basket cell. (**B, B1-B3**) Arcade axon cell. (**C, C1-C3**) Bitufted cell. Left row, the total arborization. Dashed lines indicate the border of the slice (former pial surface). Soma in gray, dendrites in black, axon in pink. Middle row, a close-up of the arborization around the parent soma. The initial axon with its collaterals highlighted in greenish color. Right row, the region boxed in the reconstructions at higher magnification to show details of the terminal endings. The axograms (A3, B3, C3) with 100 µm bin width (the thin horizontal lines) are arranged below. The axon origin is marked by the asterisks. All axograms are at the same magnification. Note the many short endings supplied by the Basket axon collaterals (A3) versus the rather long terminal elements of the arcade (B3) and bitufted cell (C3) axon collaterals.

Given the wealth of evidence that activity is promoting dendritic growth we expected to see an increase of dendritic complexity with either one of the three selected light pulse frequencies or either one of the two light pulse durations. Only the 0.5 Hz stimulation at 70 ms and 140 ms pulse duration altered the interneuronal dendrites, but unexpectedly, dendrites were on average shorter and less branched compared to the handling controls (Figure 2A and Figure 2-source data 1). The moderate decrease observed with 0.5 Hz stimulation suggested that only one of the interneuron types was affected. Since pulse duration seemed less important than pulse frequency, also in our previous study (Gonda et al., 2023), we pooled cells of the 70 ms and 140 ms condition for all following analyses. When analyzing BC (pooled) and non-BC (pooled) separately, we found that the 0.5 Hz stimulation evokes shorter and less branched dendrites selectively of non-BC (Figure 2B and Figure 2-source data 1).

**Figure 2 with Figure 2-source data 1.**
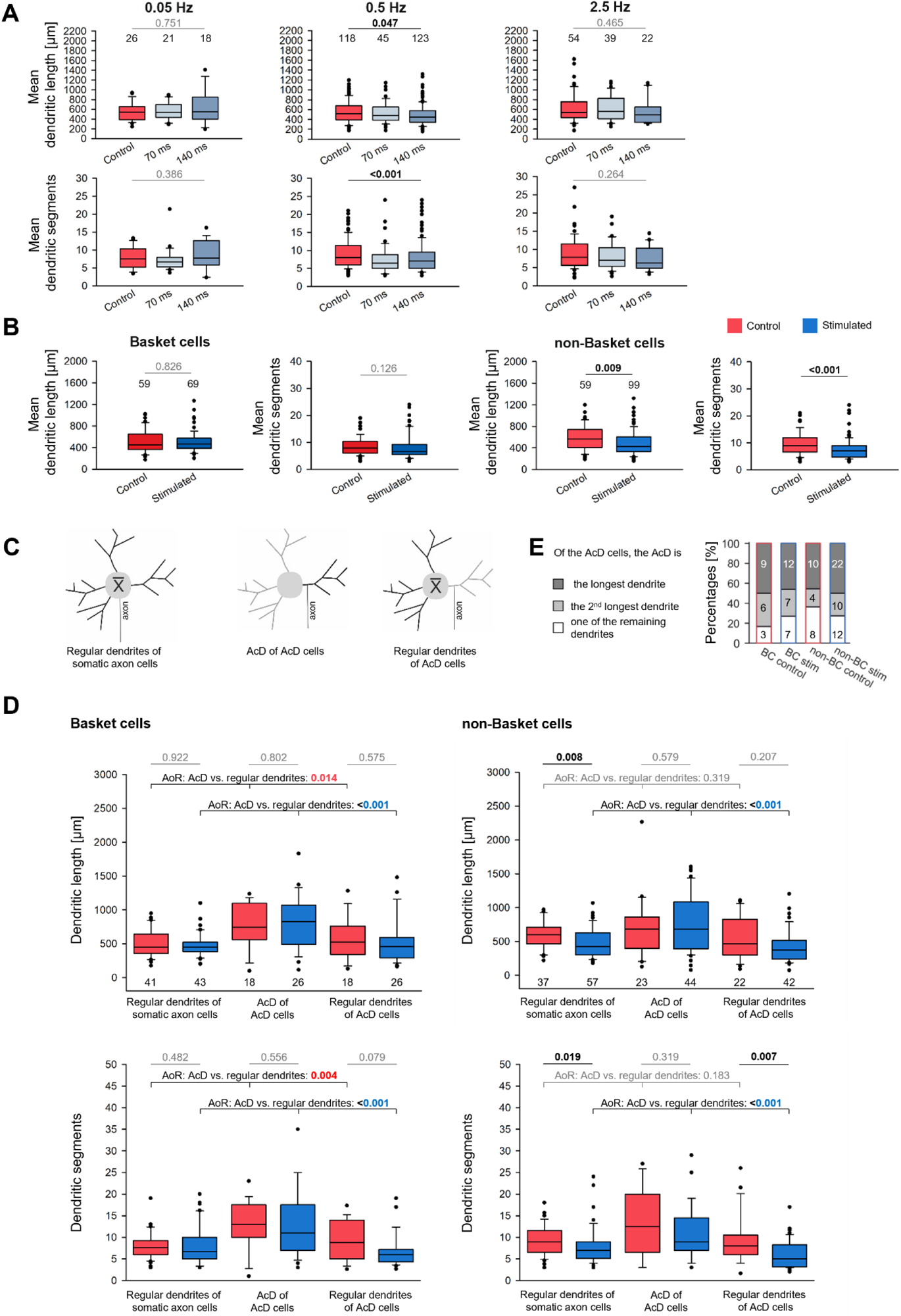
Dendrites of interneurons. (**A**) Box plots of mean dendritic length (top row) and mean dendritic segment numbers (bottom row) of all interneurons after DIV 11-15 ChR2-eYFP stimulation with 0.05 Hz, 0.5 Hz, and 2.5 Hz, each with pulse duration of 70 and 140 ms (in shades of blue). The number above each box is the number of neurons analyzed. P values determined with ANOVA on ranks corrected for multiple testing versus the handling control (in red). (**B**) Box plots of mean dendritic length (left) and mean dendritic segment numbers (right) of BC and non-BC. Handling control in red, stimulation with 70 and 140 ms conditions now pooled in blue. The number above the boxes is the number of neurons analyzed. Mann-Whitney rank sum test versus the handling control. (**C**) Sketches depicting the measured dendrites (black). The AcD of AcD cells (middle) was compared to the average of all other dendrites of the AcD cell (right), and to the average of the dendrites of non-AcD cells (left). (**D**) Length (top row) and segments (bottom row) of the three groups of dendrites for Basket cells (left column) and non-Basket cells (right column). The number of cells is given below every box. Naturally, the number of somatic axon cells is higher than the number of AcD cells. A total of 128 BC and 158 non-BC was analyzed. Of the latter, 3 cells had two axons from two dendrites, and these AcDs were included. First, we compared control to stimulated cells; Mann-Whitney rank sum test; p values are the numbers above the lines. Second, we compared the AcD versus the regular dendrites (control cells in red, 0.5 Hz stimulated cells in blue); P values determined with ANOVA on ranks (AoR). Significant differences in black or color, all other p values in gray lettering. (**E**) Percentage of BC and non-BC cells in which the AcD is the longest, second longest, or one of the remaining dendrites. **Figure 2-source data 1** Length [µm] and segment number of all interneurons reconstructed from control and optogenetically stimulated cultures @0.05 Hz, 0.5 Hz, and 2.5 Hz. Length [µm] and segment number of BC and non-BC reconstructed from control and optogenetically stimulated cultures; 0.5 Hz condition, 70 ms and 140 ms pooled. Length [µm] and segment number of AcD BC and AcD non-BC compared to regular dendrites of AcD cells and of somatic axon cells; 0.5 Hz condition, 70 ms and 140 ms pooled.

The finding mirrored our recent observation of stunted apical dendritic growth of pyramidal neurons assessed from the very same set of OTC. Also here, the effect has been only observed at 0.5 Hz (Gonda et al., 2023). The 0.05 Hz stimulation has been not effective, and the 2.5 Hz stimulation elicits dendritic injury and cell death. We suggested (Gonda et al., 2023) that the 0.05 Hz failed because it resembles the activity level typical for perinatal neurons, whereas calcium event frequencies of 2.5 Hz do not occur in visual cortex *in vivo* until well after postnatal day 20 (Rochefort et al., 2009). This might explain why pyramidal cells and interneurons during the second postnatal week respond with morphological changes only to 0.5 Hz, which comes closest to a frequency range the neurons are going to reach in their next developmental phase.

Substantial numbers of axons emerge from dendrites, termed AcDs and AcD cells, respectively. Hippocampal AcD pyramidal cells are preferentially recruited during ripple oscillations, and the AcDs display a higher excitability and an ability to elicit action potential firing which escapes somatic inhibition (Thome et al., 2014; Hodapp et al., 2022). Further, the basal AcD of CA1 pyramidal cells of 5-10 week-old mice are longer than regular dendrites and comprise about 1/3 of total basal dendritic length (Hodapp et al., 2022). However, thick-tufted cortical layer V pyramidal neurons with a basal AcD display a reduced dendritic complexity and thinner main apical dendrite than those with somatic axons (Hamada et al., 2016). We therefore analyzed the AcD versus the average of the regular dendrites as depicted in the sketch (Figure 2C and Figure 2-source data 1). For BC, AcDs of both the control cells and the 0.5 Hz stimulated cells were significantly longer and more branched than the average dendrite of somatic axon cells (left column in Figure 2D, left column). The AcDs were not extremely long, instead, they remained within the range of the longest dendrites of all individual BC of our sample.

For the non-BC of the control condition (right column in Figure 2D and Figure 2-source data 1,), the AcD were within the range of the regular dendrites. The stimulated cells with somatic axons presented on average shorter and less branched dendrites as described above (Figure 2B). The regular dendrites of the stimulated AcD cells had the lowest number of segments (right column in Figure 2D and Figure 2-source data 1). Analyzing the AcD cells separately confirmed that the AcD represented the longest or second-longest dendrite in 60-80% of the cells (Figure 2E). Together, this suggested that in BC, the AcD configuration awards a morphogenetic advantage for the dendrite which carries the axon. Further, the AcD of stimulated non-BC interneurons remained as long as the AcD of control cells. This suggested that the AcD configuration awards resistance to the growth-impairing effect of the optogenetic stimulation which stunted the regular dendrites of the AcD cells, also stimulated pyramidal cells reconstructed from the same OTCs displayed shorter apical dendrites, and infragranular cells were more susceptible than supragranular cells (Gonda et al., 2023).

### Cell type-specific effects on axonal complexity

We reconstructed the axons of BC and non-BC (70 ms and 140 ms conditions pooled). For BC and non-BC (Figure 3A, Figure 3B and Figure 3 source data 1) the total axonal length, the number of branch points of collaterals (nodes) per 1,000 µm, the maximum branch order, and the mean length of the terminal segments were not different from the handling control. Only the number of bouton terminaux tended to be higher in the stimulated non-BC axons. Comparing the two cell classes revealed differences which were to be expected. For instance, BC axons were on average somewhat longer than axons of non-BC, and BC axons had higher numbers of nodes, more bouton terminaux, and a higher maximum branch order. Quite typically, the terminal segments of the BC axon collaterals were substantially shorter and far more uniform in length than the terminal segments of non-BC axon collaterals. BC axons of comparable dimensions have been reported for P19-21 mouse dentate gyrus (Doischer et al., 2008). For non-BC, reported values in mouse cortex at P21 (Lim et al., 2018) were slightly higher than our values which might be ascribed to their slightly older age. Our axons at DIV 15 were still actively growing, yet our BC and non-BC axon dimension matched well with the data set obtained from P19 rat frontal cortex (Karube et al., 2004). Therefore, interneuronal axons in OTC are at dimensions typical for their cell type and developmental stage.

**Figure 3 with Figure 3-source data 1.**
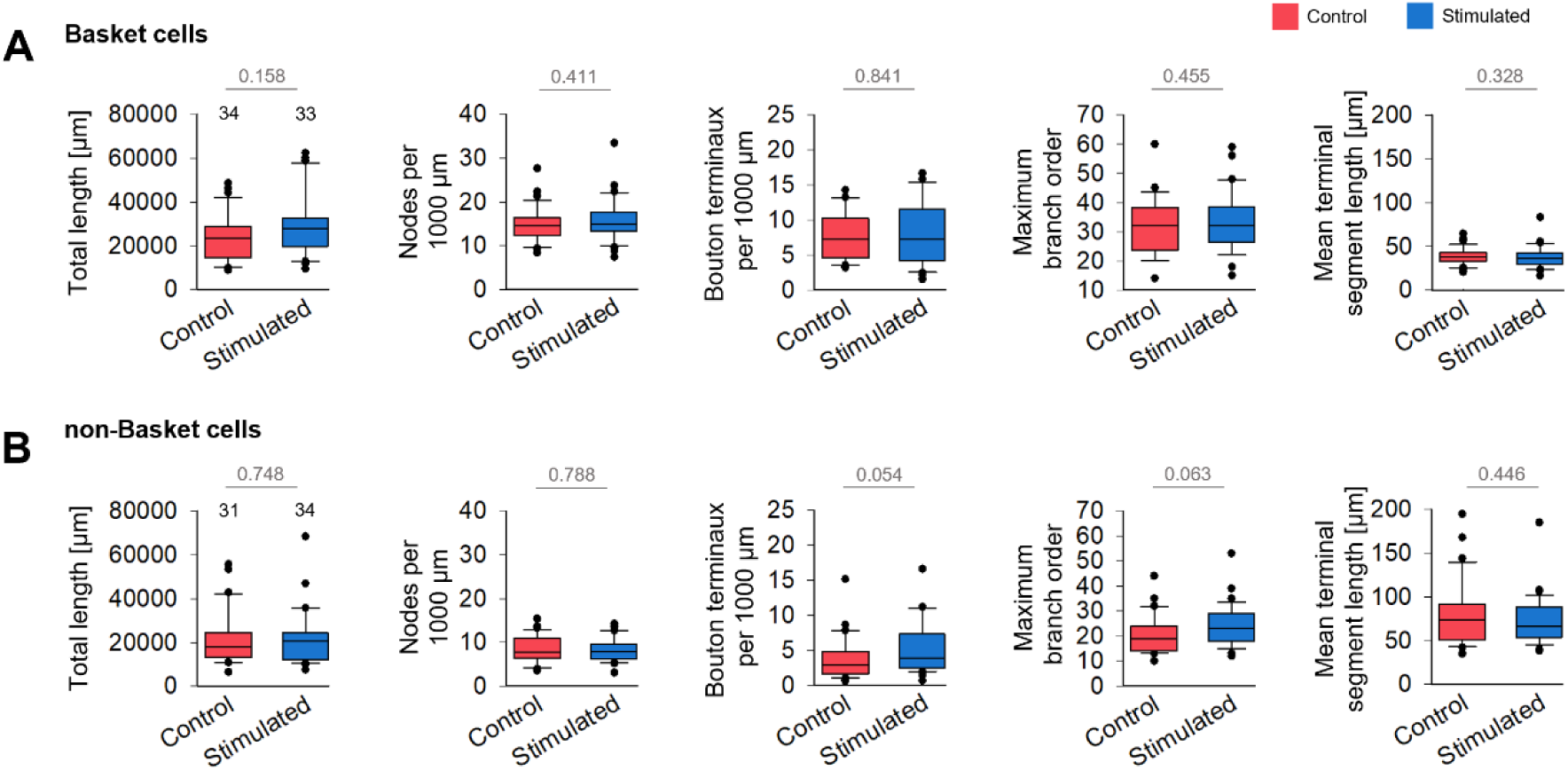
Axons of interneurons. (**A**) For BC axons, total length, number of nodes, number of bouton terminaux, maximum branch order and terminal segment length were not influenced by the 0.5 Hz stimulation. Mann-Whitney rank sum test; p values are the numbers above the lines. (**B**) For non-BC axons, total length, number of nodes, number of bouton terminaux, maximum branch order and terminal segment length were not influenced by the 0.5 Hz stimulation. Mann-Whitney rank sum test; p values are the numbers above the lines. The number above the boxes is the number of neurons analyzed. **Figure 3-source data 1** Axonal measures of BC and non-BC.

Next, Sholl-type analyses were performed (Figure 4 with Figure supplement 1 and Figure 4-source data 1). A soma-centered Sholl analysis revealed that the 0.5 Hz stimulated BC had overall more axonal intersections, and significantly more intersections within the 100-300 µm radius from the soma (Figure 4A, left). The difference became even clearer with the axogram analysis. Again, the stimulated BC had significantly more axonal intersections within the 300-600 µm linear distance from the axon origin (Figure 4B, left). In line with this, BC axons had significantly more terminal endings within the 200-400 µm bins (Figure 4C, left). No differences were seen for the non-BC axons neither in total intersection (Figure 4A, right), nor with the axogram analysis (Figure 4B, right), nor with the terminal endings (Figure 4C, right). Since the non-BC axons were reconstructed from same set of OTC, often enough from the very same culture, they represent a perfect internal control population which responded with dendritic but not axonal changes. Together, the analysis suggests that optogenetically stimulated BC axons form denser plexuses within and around the dendritic field of the parent soma.

**Figure 4 with Figure 4 supplement 1 and Figure 4-source data 1.**
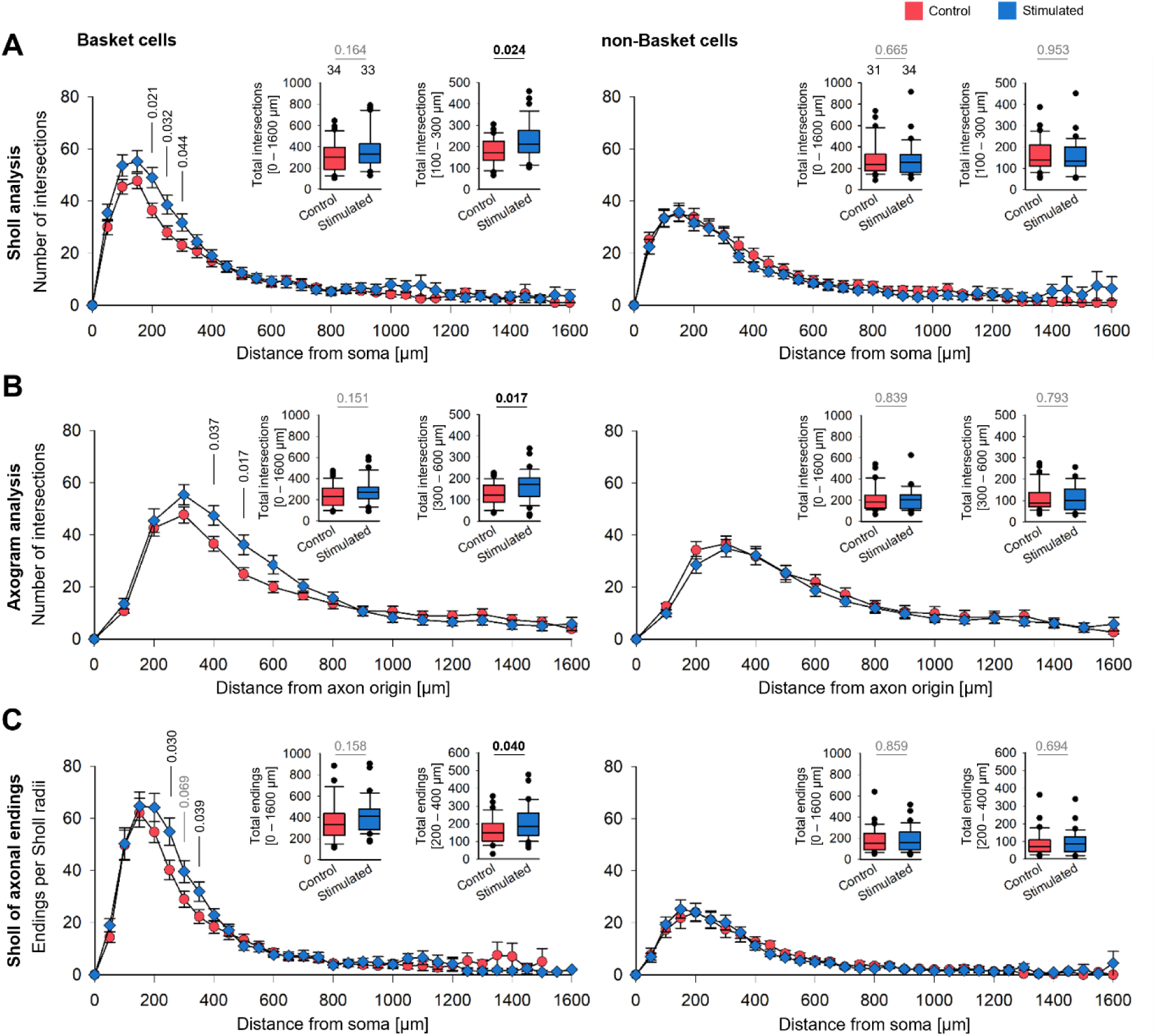
Sholl-type analyses for BC and non-BC axons. (**A**) Soma-centered Sholl analysis. (**B**) Axogram analysis. (**C**) Collateral endings per soma-centered Sholl radii. The insets depict the total intersections over the entire length of the axon (limited to 1,600 µm, a length which is not reached by all axons) and the number of intersections at selected distances from soma, resp. the axon origin (plotted as mean ± s.e.m.). Note that the bins with significant effects are shifted in the axogram analysis since it is based on the linear distance from the start point whereas the soma-centered Sholl analysis scores line crossings of collaterals weaving back and forth. P values determined by Mann-Whitney rank sum tests. Significant differences in black, all other p values in gray lettering. Handling control in red, stimulated axons in blue.

**Figure 4-Figure 4 supplement 1.**
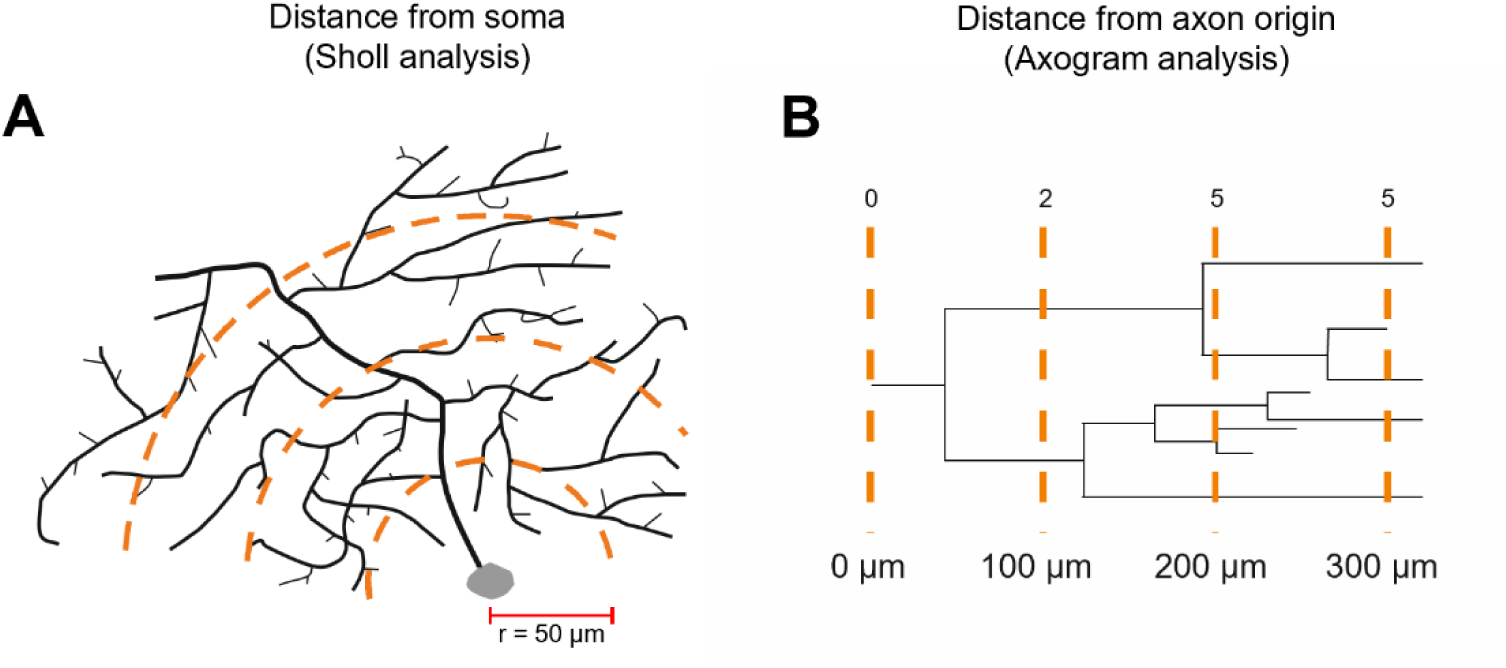
**(A)** Principle of soma-centered Sholl analysis counting crossings of circle lines at 50 µm bins. (**B**) Principle of linear axogram analysis at 100 µm bins starting at the beginning of the axon (soma or dendrite). Line crossings were determined with the MicrobrightField NeuroExplorer and Neurolucida Suite 360. **Figure 4-source data 1** Soma-centered Sholl analysis of control and stimulated BC and non-BC axons including total intersections. Axogram analysis of control and stimulated BC and non-BC axons including total intersections. Sholl analysis of terminal endings of control and stimulated BC and non-BC axons including total intersections.

The lack of effect for non-BC axons already argued for a basket cell-specific effect. However, roller cultures flatten over time in an individual manner. Therefore, we had to rule out that variable degrees of flattening of individual OTC, or portions of them, resulted in sampling errors that might have led to the statistical differences seen for BC axons (Figure 5 with Figure 5-source data 1). Any difference in culture thickness should have affected both, BC axons and non-BC axons. We thus analyzed the z-span for every cell after a 90° rotation (Figure 5A, 5B), but no difference in z-spans was observed. Also, plotting the intersections at the (arbitrarily selected) 200 µm radius of the soma-centered Sholl versus the z-span yielded no conspicuous correlations as indicated by the low r^2^ values, neither for BC (Figure 4C) nor non-BC axons (Figure 5D). As expected, the 0.5 Hz stimulated BC axons had shifted slightly upwards indicating a higher number of intersections per unit z-span (Figure 5C). This supported our finding of a stimulus-evoked increase in complexity specifically of the BC axon plexuses (Figure 4).

**Figure 5 with Figure 5-source data 1.**
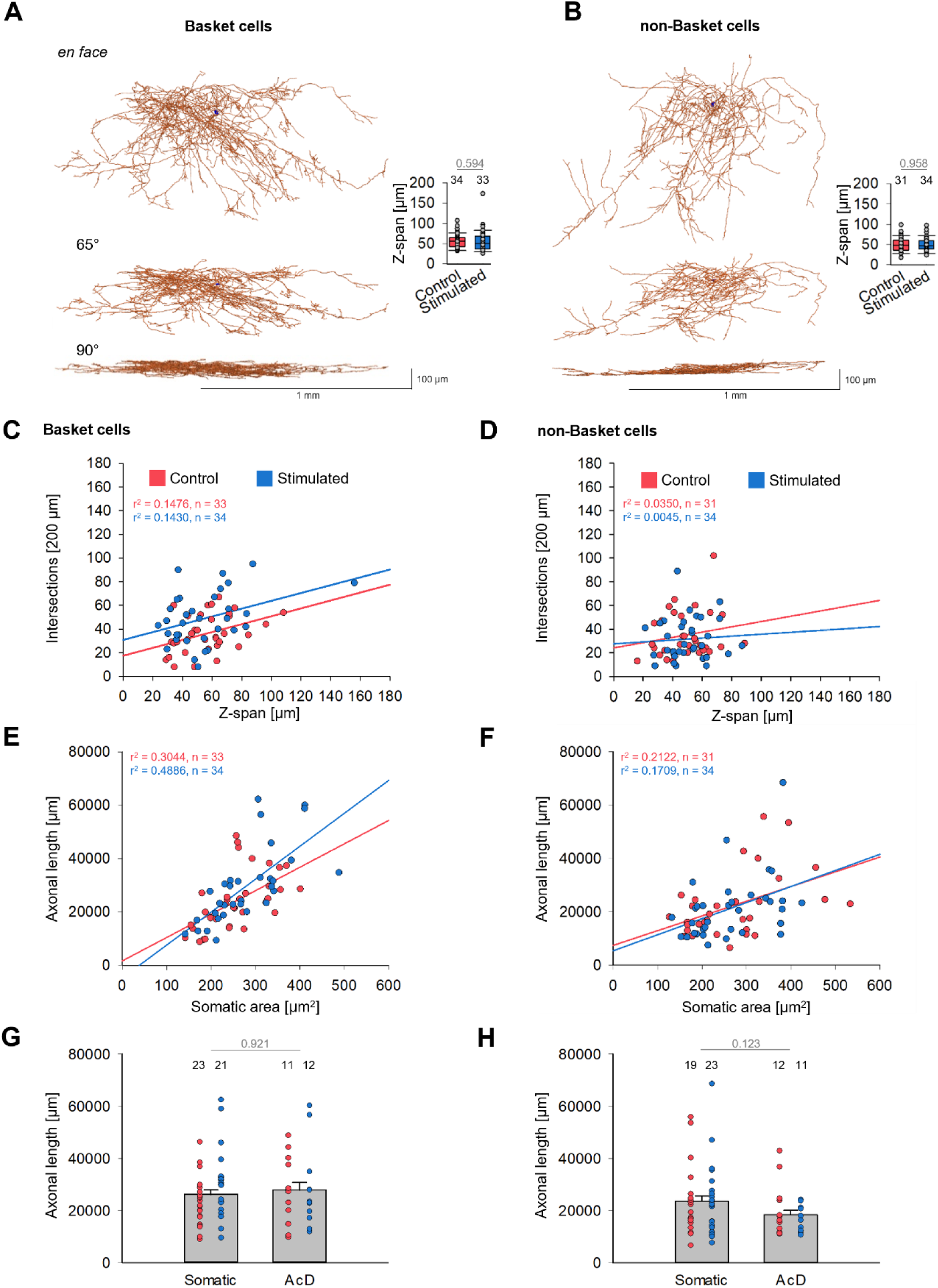
Dependence on z-span and soma size. (**A, B**) Representative BC (**A**) and non-BC (**B**) axon plexuses shown *en face* and rotated; somata in black. Insets show that control and 0.5 Hz stimulated BC and non-BC cells had on average the same z-span within the slice cultures. The number above the boxes is the number of neurons analyzed. P values determined with Mann-Whitney rank sum test. (**C, D**) Plots of Sholl circle intersections at 200 µm distance from the soma versus the z-span for BC (**C**) and non-BC (**D**). Every dot represents one neuron. (**E, F**) Plots of axon length versus soma size for BC (**E**) and non-BC (**F**) cells reveal somewhat stronger correlation in particular for the BC. Every dot is one neuron. The regression lines were computed with SigmaPlot12. (**G, H**) Length of somatic and AcD axons (gray bars; mean ± s.e.m.), and upon separation by axonal origin (red and blue dots; every dot is one cells). The numbers above the symbols are the number of axons analyzed. **Figure 5-source data 1** Sholl circle intersections at 200 µm radius versus the z-span of the axonal plexus for BC and non-BC. Total axonal length versus z-span of the plexus, and versus somatic area of BC and non-BC. Axonal length analysis with regard to axonal origin.

Interneurons differ in soma size, and large somata often support long axons. Analyzing axon length versus somatic area yielded a positive correlation for the stimulated BC cells (Figure 5E) but not for non-BC cells (Figure 5F). However, there may be no causal dependence, and we expected a certain correlation because it is well known that soma size itself is regulated by activity. Further, the axonal length was not correlated with axon origin (Figure 5G, 5H). Finally, for a set of AcD cells, the distance between soma and axon origin was determined. The distance for BC ranged from 4.9 µm to 18.8 µm (average 9.8 µm; n = 13 arbitrarily selected BC) and for non-BC the distance ranged from 1.8 µm to 32.4 µm (average 12.2 µm; n = 34).

### A morphogenetic role for the AcD of Basket neurons

The AcD has been identified as belonging to the larger dendrites of BC and non-BC (Figure 2D, 2E). When axons arise from a dendrite the two neurites share a joint segment. At the molecular level, dendrites and axons are characterized by different structural components. Competition for building material and transportation chains through the needle eye of the joint segment might put one of the neurites at a disadvantage. Thus, the axon from that dendrite might be shorter or less branched compared to axons of somatic origin. On the other hand, the AcD is a dendrite with higher excitability as determined for CA1 pyramidal cells (Thome et al., 2014; Hodapp et al., 2022). This may promote axonal development. We therefore reanalyzed the BC and non-BC sample separating cells with AcD and with somatic origin (Figure 6 and Figure 6-source data 1).

**Figure 6 with Figure 6-source data 1.**
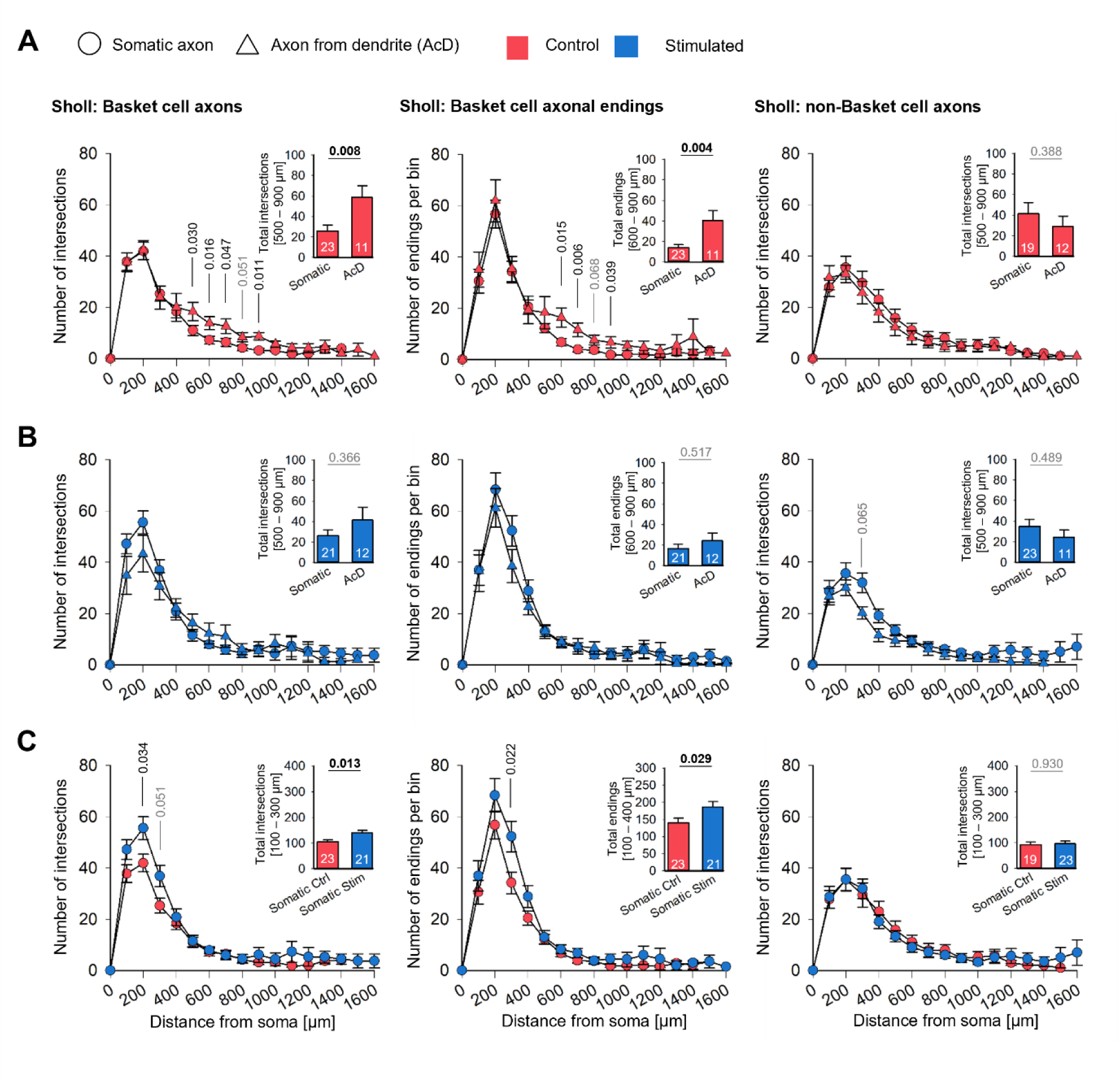
Sholl-type analyses for BC and non-BC axons originating from soma or dendrite. (**A**) Comparison of control axons originating from somata or from dendrites. (**B**) Comparison of 0.5 Hz stimulated axons originating from somata or from dendrites. (**C**) Comparison of control and stimulated axons originating from somata. Left column, intersections of BC axons; middle column, axonal endings of BC axons; right column, intersections of non-BC axons. The insets show the total intersections in the selected bin (mean ± s.e.m). Numbers in the bars are the number of axons analyzed. P values determined by Mann-Whitney rank sum tests. Significant differences in black, all other p values in gray lettering. **Figure 6-source data 1** Handling control axons separated by origin from soma or dendrite. The 0.5 Hz stimulated axons separated by origin from soma or dendrite. Handling control and stimulated axons originating from somata.

Of the control BC, axons originating from a dendrite had significantly more intersections within 500-900 µm distance from the soma than axons originating from somata (Figure 6A, left). Accordingly, the number of terminals was higher (Figure 6A, middle). No such difference was seen for non-BC axons (Figure 6A, right).

Comparing intersections and terminal endings of the optogenetically stimulated BC revealed that axons originating from somata were locally as dense as axons originating from an AcD (Figure 6B, left and 6B middle). For stimulated non-BC axons, the curves were not different (Figure 4B, right). The effect became clear when comparing axons with somatic origin of control BC to those of stimulated BC: The stimulated axons had significantly more intersections within 100-300 µm distance to the soma (Figure 6C, right) and more terminals within 100-400 µm distance to the soma (Figure 6C, middle). Again, non-BC axons originating from an AcD were not different from somatic axons, and the optogenetic stimulation did not increase the local complexity (Figure 6C, right).

Together, the results suggested that BC axons originating from a dendrite form denser local arborizations with more terminal endings within and around the dendritic field of their parent somata already in the absence of an optogenetic stimulation. Thus, morphogenesis has been driven by spontaneous activity. Optogenetic stimulation then enabled BC with somatic axons to reach the same degree of local arborization within the time given.

## DISCUSSION

The emergence of axons from dendrites of mammalian cortical neurons has been known for decades. Also, for decades, the functional implications have not been recognized. Recent work demonstrates that this neuromorphological feature is neither rare nor random. Rather, it segregates with species and cell types in the neocortex (Wahle et al., 2022). About 25% of the BC and up to 50% of the non-BC in cortex *in vivo* and in slice cultures have an axon originating from a dendrite (Höfflin et al., 2017; Wahle et al., 2022). This naturally occurring feature now delivers strong support for the view that the growth of BC dendrites and complexity of BC axons is driven by spontaneous activity. In line with our finding, the AcDs of hippocampal pyramidal cells are longer than regular dendrites of cells with somatic axons, and further, the hippocampal AcDs have a higher excitability and can elicit action potential firing which escapes somatic inhibition (Thome et al., 2014; Hodapp et al., 2022). Another example for AcDs being special dendrites comes from a recent study on chick tectal Shepherd’s crook neurons (Weigel et al., 2022). These neurons typically possess an apical AcD which receives excitatory visual input whereas the basal dendrite receives auditory input. Synaptic activation of the apical dendrites was slower and evoked more prolonged currents due to a dendritic-specific contribution of NMDA receptors (Weigel et al., 2022). Although it remains to be proven, it is tempting to speculate that the AcD of neocortical BC share similar features. The privilege of short-circuit action potential generation appears to be beneficial for both, AcD and axon of BC. Branching and terminal density increase within and around the dendritic field of the parent AcD cell. Suggestively, this may help AcD axons to form synaptic connections more rapidly and deliver more inhibition to nearby pyramidal cells from which the parent BC receives most of its excitatory input.

The AcD and its axon may mutually support each other. One might have assumed that metabolic or material transport via the needle eye of the joint segment becomes limited but at least in the time window analyzed this apparently did not delay the growth of either neurite. BC richly express calcium permeable AMPA receptors, excitatory synapses onto BC dendrites are strong, and interneuronal dendritic tip elongation comes with the formation of new excitatory synapses (Chen et al., 2011). Possibly, action potential backpropagation might contribute to stabilize the larger dendrite. NMDA receptor signaling has been shown in developing mouse olfactory bulb mitral cells to stabilize one of the primary dendrites by suppressing RhoA, and concurrently this “winner” dendrite triggers the pruning of the other dendrites (Fujimoto et al., 2023) Further, parvalbumin interneurons richly express tropomyosin receptor kinase B (TrkB) (Gorba and Wahle 1999). The dendritogenetic effect of axonal TrkB receptor activation and axonal signaling endosomes has been demonstrated for cortical neurons (Moya-Alvarado et al., 2023). The AcD is at an ideal position for receiving a major share of endosomal signals. However, there seem to be limits because the optogenetic stimulation was not able to increase the AcD’s local complexity further. By contrast, in BC with somatic axons, a comparable axonal complexity has been evoked with the optogenetic stimulation. It remains to be shown if these more complex basket cell axons elicit a stronger inhibition. As hypothetized (Gonda et al., 2023), a stronger inhibition may have contributed to the stunted growth of pyramidal cells in optogenetically stimulated OTC.

Cortical interneuron types may have distinct developmental periods during which they are most sensitive to activation or deprivation. Thus, the growth-promoting effect which we demonstrated to be specific for BC might have been due to the analyzed time window, which overlaps best with BC differentiation. Axo-somatic innervation begins to form between P5-P9, axo-somatic presynapses are well developed at P14, and mIPSC frequency increases from P6-P15 in mouse somatosensory cortex pyramidal cells with most IPSC originating from perisomatic innervation (Kobayashi et al 2008; Micheva et al., 2021). In line with this, exposure to the pro-inflammatory cytokine leukemia inhibiting factor from DIV12 onwards impairs the morphological and neurochemical differentiation of BC, whereas non-BC are barely affected (Engelhardt et al., 2018). Silencing the excitatory synaptic inputs in juvenile mice impairs dentate gyrus BC axon structure and granule cell inhibition due to subnormal NMDA receptor signaling in the BC (Pieraut et al., 2014; Feng et al., 2021). It remains to be shown if the feature “complex AcD/complex axon” persists through older ages or if axons of somatic origin will catch up in dimensions given some more time. To this end the results suggest that the AcD configuration offers a developmental advantage to BC.

For the non-BC, the lack of any axonal effect was surprising when considering that non-BC have AcDs to a much higher proportion of up to 50%, and many axons emerge with substantial distance to the soma (Höfflin et al., 2017; Wahle et al., 2022). The dendrite-targeting axonal innervation of pyramidal cells develops earlier than axo-somatic innervation starting at P5 in mouse somatosensory cortex (Gour et al., 2021). Reducing non-BC excitability by expression of Kir2.1 channels from E15 in mouse cortex drastically decreases the average axonal length of calretinin and reelin neurons at P8 (De Marco Garcia et al., 2011). Possibly, the AcD of non-BC has a growth promoting effect at younger ages. Yet, the non-BC were responding to the DIV 11-15 optogenetic stimulation, albeit with a reduction of dendritic growth. Similarly, we recently reported that also optogenetically stimulated pyramidal neurons analyzed from the same culture material develop shorter dendrites (Gonda et al., 2023). Interneurons have been shown to constantly remodel their distal dendrites (Chen et al., 2011). Such a dynamic might render them sensitive to activity-dependent regressive events. However, the AcD of the non-BC was spared and it remained among the longest dendrites suggestive of a positive influence of the AcD configuration in non-BC. To this end, we present the first evidence for a morphogenetic role of the AcD configuration.

## Materials and methods

### Culture preparation, transfection

Interneuronal axons and dendrites were reconstructed from the very same cultures batches used for analysis of pyramidal cell dendrites (Gonda et al., 2023). Briefly, roller-type cultures were prepared from postnatal day 1/2 Long-Evans hooded rat visual cortex. Parasagittal slicing anticipated the tendency of large BC to extend more in anterior-posterior direction as revealed by biocytin-filled BC axons reconstructed from stacks of 80 µm thick tangential sections of cat visual cortex covering fields of 2.3 x 2.2 and 3.8 x 1.7 mm^2^ with their radiating collaterals (Kisvarday et al., 1993). In slice cultures, the outgrowing main axonal arms will eventually adapt to the more 2D environment as roller cultures flatten over time. At DIV 8, cultures were transfected (Helios Gene Gun, Bio-Rad, Munich, Germany) with plasmids encoding CMV-driven hChR2(H134R)-eYFP (RRID: Addgene_20940).

### Optogenetic stimulation and selection of pulse frequency and duration

Cultures were maintained in a dark room to minimize exposure to white light. Stimulation was done with custom-designed illumination setup equipped with 3×8 individually switchable 465 nm LED (Osram Oslon SSL80; Lumitronix), computer-controlled via an Arduino unit as recently described in detail (Gonda et al., 2023). The distance between cultures and LED was 12 mm. Light intensity determined with a photodiode (S130VC; Thorlabs) and a power meter (PM100D (Thorlabs) at the level of the cultures in the culture tubes was 0.7 mW/mm^2^. Cultures were exposed from DIV 11-15 to daily three rounds of blue LED stimulation @0.5 Hz. Each round had 3 x 5 min stimulation with 5 min pauses during which the handling controls were mock-stimulated in the setup, a 30 min break, and again 3 x 5 min with 5 min pauses for mock-stimulating the controls; this way each round of stimulation lasted 1.5 h in total. The 0.5 Hz frequency has been effective in altering pyramidal cell dendritic growth (Gonda et al., 2023). Whole cell illumination efficiently evokes neuronal spiking and action potential backpropagation (Grossman et al., 2013).

Previous work has shown that overexpression of specific glutamate receptor subunits increases dendritic complexity because the amplitude of depolarizing events increases with more receptors (Hamad et al., 2011; Jack et al., 2019). Event frequencies in our cultures around DIV 10 are ∼0.01 Hz, and for instance the overexpression of GluK2(Q) in pyramidal cells causes an increase of the frequency of calcium events to ∼0.06 Hz which results in the growth of apical dendrites (Jack et al., 2019). Calcium events were at 0.02–0.04 Hz at DIV 11-15, and at DIV 18-20 individual pyramidal cells expressing genetically encoded calcium indicators display 0.1 Hz (Hamad et al., 2014; Jack et al., 2019; Engelhardt et al., 2018; Gasterstädt et al., 2022). The lowest frequency of 0.01 Hz matches the frequency of spontaneously occurring calcium events reported for supragranular neurons of the anesthetized mouse primary visual cortex at postnatal day 8, and at postnatal day 11 ∼0.03 Hz are reported (Rochefort et al., 2009). Calcium event frequency of ∼0.25 Hz occur after eye opening and further increase to 0.5 Hz by the end of the fourth postnatal week (Rochefort et al., 2009), while the number of cells contributing to the events decreases substantially after eye opening *in vivo* (Rochefort et al., 2009). Together, this determined our choice of 0.05 Hz as „immature” frequency, 0.5 Hz as frequency reached during the third postnatal week, and 2.5 Hz as a frequency which is not naturally occuring during our window of analysis. In fact, the 0.5 Hz stimulation turned out effective, whereas the 0.05 Hz frequency was not altering dendritic growth and the 2.5 Hz evoked dendritic injury (Gonda et al., 2023). Light pulse duration was 70 ms and 140 ms. In particular the latter evoked a strong increase of c-Fos protein in ChR2-transfected neurons, and both durations were altering pyramidal cell dendritic growth. Therefore, as done previously (Gonda et al., 2023) we pooled the neurons of the 70 ms and 140 ms conditions for the axon analysis.

### Immunostaining

At DIV 15, ∼3 h after the final round of stimulation, cultures were fixed and immunostained with mouse anti-GFP antibody (1:1000; clone GSN24, Sigma-Aldrich, RRID: AB_563117) as described (Gonda et al., 2020; Gasterstädt et al., 2022). The batch-internal “handling” control received the same ChR2-eYFP transfection, and the same time in the illumination setup without any LED light making the stimulation the only experimental variable. A handling control is essential because placing culture tubes in and out of the illumination setup, as gentle as it was done, could potentially generate mechanical stress which has been shown to contribute to dendritic remodeling (Franze et al., 2009).

### Inclusion and exclusion criteria for selection of interneuron types

BC feature quite thick initial axons emerging from the soma or from a dendrite (Kisvarday et al., 1993; Höfflin et al., 2017; Wahle et al., 2022). The initial axon gives rise to major collaterals forming horizontal connections in L2/3 resp. infragranular layers. Delicate collaterals emerge which quicky become varicose and form a dense local plexus within the dendritic field. Collaterals are studded with irregular-sized and partly large boutons and numerous short varicose axo-somatic terminal elements ending in close apposition to pyramidal-shaped somata and their proximal dendrites. Therefore, by inclusion criteria our sample comprised the classical large horizontal BC, and smaller BC with more local plexuses (Figure 1 - Figure supplement 1A, A1-A3).

Of the non-Basket (non-BC) cells we included cells with arcade-shaped axon arbors (Figure 1B - Figure supplement 1B, B1-B3) and bitufted cells (Figure 1-Figure supplement 1C, C1-C3), which were most frequently labeled in our material. Non-BC are primarily dendrite-targeting cells with adapting firing pattern (Markram et al., 2015; Feldmeyer et al., 2018). Their axons often emerge from dendrites (Höfflin et al., 2017; Wahle et al., 2022), and branch more sparsely than BC axons. Collaterals have rather small regular-sized boutons and project translaminar without extending into layer 1.

We excluded Martinotti cells for their highly variable axon length in layer 1. Somatostatin-positive Martinotti cell axons have been measured after genetic labeling in mouse cortex at postnatal day 21. Total axonal length varies from ∼5 to >25 mm (Lim et al., 2018). Harvesting by chance a few Martinotti cells with extremely long or very short collaterals in layer 1 for either one of our experimental conditions could have easily led to false-positive results. Axons of translaminar somatostatin cells (types included in our study) display an average length of 25-30 mm at P21 and thus had a lower individual variability (Lim et al., 2018). Chandelier cells were not included due to immaturity, because axo-axonic innervation becomes recognizable by P14 and older (Pan-Vasquez et al., 2020; Gour et al., 2021; Micheva et al., 2021), beyond the time window analyzed here. We thus excluded layer 2/3 interneurons with prominent dendrites oriented towards layer 1, a typical feature of chandelier cells (Taniguchi et al., 2013). We also excluded bipolar cells because their small somata were too rarely transfected with biolistics and their axons are thin, sparsely branched, and not always entirely labeled. Last, we excluded occasionally transfected neurogliaform neurons recognizable by their rather smooth axons of thin caliber and short dendrites (Staiger et al., 2015; Feldmeyer et al., 2018).

### Morphometry

We reconstructed without preselection all neurons of the desired types which had completely stained dendrites and/or completely stained axonal plexues in addition. With sparse transfections, cells ideally resided in a solitary position allowing for 3D manual reconstruction (Neurolucida, MicroBrightField, Williston, USA) done by two trained observers who were blinded to condition. Reconstructions were analyzed with the Neurolucida software and the Neurolucida 360 Suite. In total, dendrites of 466 interneurons were reconstructed (n = 65 cells @0.05 Hz; 286 cells @0.5 Hz; 115 cells @2.5 Hz). For dendrites, we determined the mean dendritic length and segment number per cell for the three stimulus frequencies. Only the 0.5 Hz stimulation altered interneuronal dendritic complexity, and this was the case also for pyramidal neurons (Gonda et al., 2023). Therefore, we analyzed the 0.5 Hz sample separated by cell type (BC: 59 control cells; 69 stimulated cells; non-BC: 59 control cells; 99 stimulated cells). Because we did not see evidence for a distinct effect of stimulus duration, we pooled the reconstructed cells of the 70 ms and 140 ms conditions.

Further, from the 0.5 Hz sample, we reconstructed a total of 132 completely stained interneuronal axons (n = 67 BC axons; 65 non-BC axons; reconstructed axonal length in total ∼3.2 meter). For axons, we determined total length, the number of branch points (nodes; all classified as bifurcations) per 1,000 µm, bouton terminaux per 1,000 µm, maximum branch order, and the mean terminal segment length. Complexity of the axon and spatial distribution of the terminal endings were analysed with soma centered Sholl analysis and 1D linear axograms starting at the axon origin which may be somatic or dendritic. The z-span of the axon in the OTC was determined after 90° rotation or the reconstruction and measuring the largest distance reached by collaterals. Somatic area was determined as proxy for soma size. The Sholl analysis of axonal endings was done with 50 µm bins. Next, we averaged the counts of two adjacent bins to work against variability (terminal endings are sometimes densely clustered). We eventually plotted the average, and the graphs thus report the average number of terminal endings of every second bin, plotted at 100 µm distance to the soma. This way the axes of the graphs could have been kept at the same dimension for easier comparison between the plots.

### Statistical analysis

Reconstructions derive from 86 slice cultures from 19 independent preparations, each done with 5-6 perinatal rats of both sexes, and slices from every animal were allocated to all experimental conditions run with each preparation. Graphs and statistics were done with SigmaStat12.3 (Systat Software GmbH, Frankfurt am Main, Germany). Statistical comparison was done using non-parametric Mann-Whitney rank sum tests or ANOVA on ranks with Dunn’s correction for multiple testing where appropriate. All data presented in the graphs are provided in the first data sheet of the Excel source data files; detailed information on statistical comparisons showing a significant difference is given in the second data sheet of the Excel source data files to Figure 2, Figure 4, and Figure 6.

## Acknowledgments

We thank Andrea Räk and Sabine Kleinhubbert for technical support.

## Author contributions

Steffen Gonda, Methodology, Investigation, Formal analysis, Visualization; Christian Riedel, Methodology, Investigation; Andreas Reiner, Methodology, Writing – original draft; Ina Köhler, Methodology, Investigation, Formal analysis; Petra Wahle, Conceptualization, Formal analysis, Funding acquisition, Investigation, Methodology, Project administration, Resources, Supervision, Visualization, Writing - original draft, Writing – review and editing

## Author ORCIDs

Steffen Gonda http://orcid.org/0000-0003-1605-8605

Christian Riedel http://orcid.org/0000-0001-9998-1563

Andreas Reiner http://orcid.org/0000-0003-0802-7278

Ina Köhler http://orcid.org/0000-0002-0709-9598

Petra Wahle http://orcid.org/0000-0002-8710-0375

## Ethics

Work was done in compliance with Ruhr University Bochum Animal Research Board and the Federal State of North Rhine-Westphalia.

## Data and materials availability

All data generated or analysed during this study are included in the manuscript and supporting file; source data files have been provided for Figures 2, 3, 4, 5, and 6.

## REFERENCES

Alhourani A, Fish KN, Wozny TA, Sudhakar V, Hamilton RL, Richardson RM. 2020. GABA bouton subpopulations in the human dentate gyrus are differentially altered in mesial temporal lobe epilepsy. The Journal of Neurophysiology 123:392–406. 10.1152/jn.00523.2018, PMID: 31800363

Amegandjin CA, Choudhury M, Jadhav V, Carriço JN, Quintal A, Berryer M, Snapyan M, Chattopadhyaya B, Saghatelyan A, Di Cristo G. 2021. Sensitive period for rescuing parvalbumin interneurons connectivity and social behavior deficits caused by TSC1 loss. Nature Communications 12:3653. 10.1038/s41467-021-23939-7, PMID: 34135323

Baho E, Di Cristo G. 2012. Neural activity and neurotransmission regulate the maturation of the innervation field of cortical GABAergic interneurons in an age-dependent manner. The Journal of Neuroscience 32:911–918. 10.1523/JNEUROSCI.4352-11.2012, PMID: 22262889

Chattopadhyaya B, Di Cristo G, Wu CZ, Knott G, Kuhlman S, Fu Y, Palmiter RD, Huang ZJ. 2007. GAD67-mediated GABA synthesis and signaling regulate inhibitory synaptic innervation in the visual cortex. Neuron 54:889–903. 10.1016/j.neuron.2007.05.015, PMID: 17582330

Chattopadhyaya B, Di Cristo G, Higashiyama H, Knott GW, Kuhlman SJ, Welker E, Huang ZJ. 2004. Experience and activity-dependent maturation of perisomatic GABAergic innervation in primary visual cortex during a postnatal critical period. The Journal of Neuroscience 24:9598–9611. 10.1523/JNEUROSCI.1851-04.2004, PMID: 15509747

Chen JL, Lin WC, Cha JW, So PT, Kubota Y, Nedivi E. 2011. Structural basis for the role of inhibition in facilitating adult brain plasticity. Nature Neuroscience 14:587–594. 10.1038/nn.2799, PMID: 21478885

De Marco García NV, Karayannis T, Fishell G. 2011. Neuronal activity is required for the development of specific cortical interneuron subtypes. Nature 472:351–355. 10.1038/nature09865, PMID: 21460837

Di Cristo G, Wu C, Chattopadhyaya B, Ango F, Knott G, Welker E, Svoboda K, Huang ZJ. 2004. Subcellular domain-restricted GABAergic innervation in primary visual cortex in the absence of sensory and thalamic inputs. Nature Neuroscience 7:1184–1186. 10.1038/nn1334, PMID: 15475951

Doischer D, Hosp JA, Yanagawa Y, Obata K, Jonas P, Vida I, Bartos M. 2008. Postnatal differentiation of basket cells from slow to fast signaling devices. The Journal of Neuroscience 28:12956–12968. 10.1523/JNEUROSCI.2890-08.2008, PMID: 19036989

Engelhardt M, Hamad MIK, Jack A, Ahmed K, König J, Rennau LM, Jamann N, Räk A, Schönfelder S, Riedel C, Wirth MJ, Patz S, Wahle P. 2018. Interneuron synaptopathy in developing rat cortex induced by the pro-inflammatory cytokine LIF. Experimental Neurology 302:169–180. 10.1016/j.expneurol.2017.12.011, PMID: 29305051

Favuzzi E, Deogracias R, Marques-Smith A, Maeso P, Jezequel J, Exposito-Alonso D, Balia M, Kroon T, Hinojosa AJ, F Maraver E, Rico B. 2019. Distinct molecular programs regulate synapse specificity in cortical inhibitory circuits. Science 363:413–417. 10.1126/science.aau8977, PMID: 30679375

Feldmeyer D, Qi G, Emmenegger V, Staiger JF. 2018. Inhibitory interneurons and their circuit motifs in the many layers of the barrel cortex. Neuroscience 368:132–151. 10.1016/j.neuroscience.2017.05.027, PMID: 28528964

Feng T, Alicea C, Pham V, Kirk A, Pieraut S. 2021. Experience-dependent inhibitory plasticity is mediated by CCK+ basket cells in the developing dentate gyrus. The Journal of Neuroscience 41:4607–4619. 10.1523/JNEUROSCI.1207-20.2021, PMID: 33906898

Franze K, Gerdelmann J, Weick M, Betz T, Pawlizak S, Lakadamyali M., et al. 2009. Neurite branch retraction is caused by a threshold-dependent mechanical impact. Biophysical Journal 97:1883–1890. 10.1016/j.bpj.2009.07.033, PMID: 19804718

Fujimoto S, Leiwe MN, Aihara S, Sakaguchi R, Muroyama Y, Kobayakawa R, Kobayakawa K, Saito T, Imai T. 2023. Activity-dependent local protection and lateral inhibition control synaptic competition in developing mitral cells in mice. Developmental Cell 58:1221–1236.e7. 10.1016/j.devcel.2023.05.004, PMID: 37290446

Gasterstädt I, Schröder M, Cronin L, Kusch J, Rennau LM, Mücher B, Herlitze S, Jack A, Wahle P. 2022. Chemogenetic silencing of differentiating cortical neurons impairs dendritic and axonal growth. Frontiers in Cellular Neuroscence. 16, 941620. 10.3389/fncel.2022.941620 PMID: 35910251

Gonda S, Giesen J, Sieberath A, West F, Buchholz R, Klatt O, Ziebarth T, Räk A, Kleinhubbert S, Riedel C, Hollmann M, Hamad MIK, Reiner A, Wahle P. 2020. GluN2B but not GluN2A for basal dendritic growth of cortical pyramidal neurons. Frontiers in Neuroanatomy 14:571351. 10.3389/fnana.2020.571351, PMID: 33281565

Gonda S, Köhler I, Haase A, Czubay K, Räk A, Riedel C, Wahle P. 2023. Optogenetic stimulation shapes dendritic trees of infragranular cortical pyramidal cells. Frontiers in Cellular Neuroscience 17: 1212483. 10.3389/fncel.2023.1212483, PMID: 37587917

Gorba T, Wahle P. 1999. Expression of TrkB and TrkC but not BDNF mRNA in neurochemically identified interneurons in rat visual cortex in vivo and in organotypic cultures. The European Journal of Neuroscience 11:1179–1190. 10.1046/j.1460-9568.1999.00551.x, PMID: 10103114

Gour A, Boergens KM, Heike N, Hua Y, Laserstein P, Song K, Helmstaedter M. 2021. Postnatal connectomic development of inhibition in mouse barrel cortex. Science 371:eabb4534. 10.1126/science.abb4534, PMID: 33273061

Grossman N, Poher V, Grubb MS, Kennedy GT, Nikolic K, McGovern B, Berlinguer Palmini R, Gong Z, Drakakis EM, Neil MA, Dawson MD, Burrone J, Degenaar P. 2010. Multi-site optical excitation using ChR2 and micro-LED array. Journal Neural Engineering 7:16004. 10.1088/1741-2560/7/1/016004, PMID: 20075504

Hamad MI, Ma-Högemeier ZL, Riedel C, Conrads C, Veitinger T, Habijan T, Schulz JN, Krause M, Wirth MJ, Hollmann M, Wahle P. 2011. Cell class-specific regulation of neocortical dendrite and spine growth by AMPA receptor splice and editing variants. Development 138:4301–4313. 10.1242/dev.071076. PMID: 21865324

Hamada MS, Goethals S, de Vries SI, Brette R, Kole MH. 2016. Covariation of axon initial segment location and dendritic tree normalizes the somatic action potential. Proceedings National Academy of Science U.S.A. 113:14841–14846. 10.1073/pnas.1607548113, PMID: 27930291

Hodapp A, Kaiser ME, Thome C, Ding L, Rozov A, Klumpp M, Stevens N, Stingl M, Sackmann T, Lehmann N, Draguhn A, Burgalossi A, Engelhardt M, Both M. 2022. Dendritic axon origin enables information gating by perisomatic inhibition in pyramidal neurons. Science 377:1448–1452. 10.1126/science.abj1861, PMID: 36137045

Höfflin F, Jack A, Riedel C, Mack-Bucher J, Roos J, Corcelli C, Schultz C, Wahle P, Engelhardt M. 2017. Heterogeneity of the axon initial segment in interneurons and pyramidal cells of rodent visual cortex. Frontiers in Cellular Neuroscience 11:332. 10.3389/fncel.2017.00332, PMID: 29170630

Jack A, Hamad MIK, Gonda S, Gralla S, Pahl S, Hollmann M, Wahle P. 2019. Development of cortical pyramidal cell and interneuronal dendrites: A role for kainate receptor subunits and NETO1. Molecular Neurobiology 56:4960–4979. 10.1007/s12035-018-1414-0. PMID: 30421168

Karube F, Kubota Y, Kawaguchi Y. 2004. Axon branching and synaptic bouton phenotypes in GABAergic nonpyramidal cell subtypes. The Journal of Neuroscience 24:2853–2865. 10.1523/JNEUROSCI.4814-03.2004, PMID: 15044524

Keck T, Scheuss V, Jacobsen RI, Wierenga CJ, Eysel UT, Bonhoeffer T, Hübener M. 2011. Loss of sensory input causes rapid structural changes of inhibitory neurons in adult mouse visual cortex. Neuron 71:869–882. 10.1016/j.neuron.2011.06.034, PMID: 21903080

Kisvárday ZF, Beaulieu C, Eysel UT. 1993. Network of GABAergic large basket cells in cat visual cortex (area 18): implication for lateral disinhibition. The Journal of Comparative Neurology 327:398–415. 10.1002/cne.903270307, PMID: 8440773

Klostermann O, Wahle P. 1999. Patterns of spontaneous activity and morphology of interneuron types in organotypic cortex and thalamus-cortex cultures. Neuroscience 92:1243–1259. 10.1016/s0306-4522(99)00009-3, PMID: 10426481

Kobayashi M, Hamada T, Kogo M, Yanagawa Y, Obata K, Kang Y. 2008. Developmental profile of GABAA-mediated synaptic transmission in pyramidal cells of the somatosensory cortex. The European Journal of Neuroscience 28:849–861. 10.1111/j.1460-9568.2008.06401.x, PMID: 18691332

Lim L, Pakan JMP, Selten MM, Marques-Smith A, Llorca A, Bae SE, Rochefort NL, Marín O. 2018. Optimization of interneuron function by direct coupling of cell migration and axonal targeting. Nature Neuroscience 21:920–931. 10.1038/s41593-018-0162-9, PMID: 29915195

Marik SA, Yamahachi H, McManus JN, Szabo G, Gilbert CD. 2010. Axonal dynamics of excitatory and inhibitory neurons in somatosensory cortex. PloS Biol 8:e1000395. 10.1371/journal.pbio.1000395, PMID: 20563307

Markram H, Muller E, Ramaswamy S, Reimann MW, Abdellah M, Sanchez CA, Ailamaki A, et al. 2015. Reconstruction and simulation of neocortical microcircuitry. Cell 163:456–492. 10.1016/j.cell.2015.09.029, PMID: 26451489

Meyer G, Ferres-Torres R. 1984. Postnatal maturation of nonpyramidal neurons in the visual cortex of the cat. The Journal of Comparative Neurology 228:226–244. 10.1002/cne.902280209, PMID: 6480914

Micheva KD, Kiraly M, Perez MM, Madison DV. 2021. Extensive structural remodeling of the axonal arbors of parvalbumin basket cells during development in mouse neocortex. The Journal of Neuroscience 41:9326–9339. 10.1523/JNEUROSCI.0871-21.2021, PMID: 34583957

Moya-Alvarado G, Tiburcio-Felix R, Ibáñez MR, Aguirre-Soto AA, Guerra MV, Wu C, Mobley WC, Perlson E, Bronfman FC. 2023. BDNF/TrkB signaling endosomes in axons coordinate CREB/mTOR activation and protein synthesis in the cell body to induce dendritic growth in cortical neurons. eLife 12:e77455. 10.7554/eLife.77455, PMID: 36826992

Munguba H, Chattopadhyaya B, Nilsson S, Carriço JN, Memic F, Oberst P, Batista-Brito R, Muñoz-Manchado AB, Wegner M, Fishell G, Di Cristo G, Hjerling-Leffler J. 2021., Postnatal Sox6 regulates synaptic function of cortical parvalbumin-expressing neurons. The Journal of Neuroscience 41:8876-8886. 10.1523/JNEUROSCI.0021-21.2021, PMID: 34503995

Pan-Vazquez A, Wefelmeyer W, Gonzalez Sabater V, Neves G, Burrone J. 2020. Activity-dependent plasticity of axo-axonic synapses at the axon initial segment. Neuron 106:265–276.e6. 10.1016/j.neuron.2020.01.037, PMID: 32109363

Patrizi A, Awad PN, Chattopadhyaya B, Li C, Di Cristo G, Fagiolini M. 2020. Accelerated hyper-maturation of parvalbumin circuits in the absence of MeCP2. Cerebral Cortex 30:256–268. 10.1093/cercor/bhz085, PMID: 31038696

Pieraut S, Gounko N, Sando R 3rd, Dang W, Rebboah E, Panda S, Madisen L, Zeng H, Maximov A. 2014. Experience-dependent remodeling of basket cell networks in the dentate gyrus. Neuron 84:107–122. 10.1016/j.neuron.2014.09.012, PMID: 25277456

Rochefort NL, Garaschuk O, Milos RI, Narushima M, Marandi N, Pichler B, Kovalchuk Y, Konnerth A. 2009. Sparsification of neuronal activity in the visual cortex at eye-opening. Proceedings National Academy of Science U.S.A. 106:15049–15054. 10.1073/pnas.0907660106, PMID: 19706480

Staiger JF, Möck M, Prönneke A, Witte M. 2015. What types of neocortical GABAergic neurons do really exist? Neuroforum 21:64–73. 10.1007/s12269-015-0012-6

Stedehouder J, Brizee D, Shpak G, Kushner SA. 2018. Activity-dependent myelination of parvalbumin interneurons mediated by axonal morphological plasticity. The Journal of Neuroscience 38:3631–3642. 10.1523/JNEUROSCI.0074-18.2018, PMID: 29507147

Steinecke A, Hozhabri E, Tapanes S, Ishino Y, Zeng H, Kamasawa N, Taniguchi H. 2017. Neocortical Chandelier cells developmentally shape axonal arbors through reorganization but establish subcellular synapse specificity without refinement. eNeuro 4.ENEURO.0057-17.2017. 10.1523/ENEURO.0057-17.2017, PMID: 28584877

Taniguchi H, Lu J, Huang ZJ. 2013. The spatial and temporal origin of chandelier cells in mouse neocortex. Science 339:70–74. 10.1126/science.1227622, PMID: 23180771

Thome C, Kelly T, Yanez A, Schultz C, Engelhardt M, Cambridge SB, Both M, Draguhn A, Beck H, Egorov AV. 2014. Axon-carrying dendrites convey privileged synaptic input in hippocampal neurons. Neuron 83:1418–1430. 10.1016/j.neuron.2014.08.013, PMID: 25199704

Wahle P, Sobierajski E, Gasterstädt I, Lehmann N, Weber S, Lübke JHR, Engelhardt M, Distler C, Meyer G. 2022. Neocortical pyramidal neurons with axons emerging from dendrites are frequent in non-primates, but rare in monkey and human. eLife 11:e76101. 10.7554/eLife.76101, PMID: 35441590

Weigel S, Künzel T, Lischka K, Huang G, Luksch H. 2022. Morphology and dendrite-specific synaptic properties of midbrain neurons shape multimodal integration. The Journal of Neuroscience 42:2614–2630. 10.1523/JNEUROSCI.1695-21.2022, PMID: 35135851

